# Behavioral profiling of hyperbaric oxygen as an intervention for chemotherapy-related functional impairments in male and female mice

**DOI:** 10.1101/2025.02.21.639572

**Authors:** Oanh T.P. Trinh, Andrew Ferns, Bethany S. Zachariah, Nathalie Sumien

## Abstract

‘Chemobrain’ or chemotherapy-related cognitive impairment (CRCI) affects up to 75% of cancer patients and survivors following chemotherapy treatments. Chemotherapy typically affects multiple domains including learning, memory, attention, executive function, and mood regulation, persisting for decades after treatment cessation and significantly diminishing cancer survivors’ quality of life. Despite its prevalence and long-term impact, effective interventions for CRCI remain limited. This study investigated the behavioral effects of HBO on mice exposed to chemotherapy drugs methotrexate (MTX) and 5-fluorouracil (5-FU). Adult male and female C57BL/6 mice received intraperitoneal injections of either saline or chemotherapy (Low-dose: MTX 37.5 mg/kg and 5-FU 50 mg/kg; High-dose: MTX 70 mg/kg and 5-FU 100 mg/kg) once a week for three weeks. Concurrently, subsets of mice underwent daily HBO (2.4 ATA, 90 minutes) five days a week for three weeks. Animals’ health was evaluated weekly, and behavioral assessment of cognitive, motor, and affective functions was conducted post-treatment. Our results showed that chemotherapy, especially at high-dose, impaired spatial memory and navigation, avoidance learning, fear discrimination, and anxiety regulation differently between males and females. HBO significantly alleviated chemotherapy-induced avoidance learning impairment in both sexes and improved coordinated running capacity in high-dose treated males. However, chemotherapy-HBO cotreatment increased spatial memory deficit in males and increased anxiety-like behaviors in females. In conclusion, even though HBO had some nuanced effects on the various domains, some reversal of CRCI effects were observed. Therefore, HBO should be further studied and considered as a potential treatment for ‘chemobrain’.

## 1. Introduction

Chemotherapy is a common treatment for various cancers, however it is associated with numerous adverse effects. One of these negative effects is chemotherapy-related cognitive impairment (CRCI), also referred to as post-chemotherapy cognitive impairment (PCCI), ‘chemo fog’, or ‘chemobrain’. Independent of cancer itself, chemotherapy negatively affects multiple cognitive domains, including executive function, decision-making, visual-spatial processing, attention, concentration, learning, and memory [1, 2]. Clinical studies indicate that CRCI affects a substantial proportion of cancer survivors, with reported incidence rates ranging from 17% to 75% [1]. However, prevalence varies by cancer type, affecting more than 50% of breast cancer survivors [3–5], nearly 70% of ovarian cancer survivors [6], 13.3% to 57% of colorectal cancer survivors [7], 52% of Hodgkin lymphoma survivors [8], and 62% of acute myeloid leukemia patients [9]. Additional estimates include 24% of lung cancer patients experiencing treatment-induced decision-making deficit [10] and approximately 51% of prostate cancer patients have an increased chance of experiencing cognitive problems within 12 months of chemotherapy exposure [11].

The duration of CRCI is highly variable, with cognitive symptoms persisting from few months to 20 years post-treatment [12–14]. Even minor neurocognitive declines may significantly impact quality of life, affecting daily functioning and imposing financial, emotional, and social burdens on cancer survivors [15–17]. Despite the growing recognition of CRCI, effective therapy options remain limited. A systematic review addressing CRCI interventions found that pharmacologic treatments, such as psychostimulants, epoetin alfa, and Ginkgo biloba, are largely ineffective [18]. Non-pharmacological approaches, such as psychotherapy and exercise therapy, have modest benefits in self-reported memory and executive functions but suffer from methodological limitations that hinder clinical recommendations [18, 19]. Furthermore, CRCI- preventative pharmacological intervention administered during chemotherapy may introduce drug-drug interactions, potentially reducing anti-cancer efficacy and increasing adverse events [20]. Similarly, exercise-based therapies, while beneficial, may be difficult to implement due to chemotherapy-induced fatigue and muscle weakness [21, 22]. Therefore, there is a critical need for a non-invasive intervention that accommodates the reduced physical capacity of patients and does not compromise chemotherapy efficacy.

Methotrexate (MTX) and 5-fluorouracil (5-FU) are widely used cytotoxic agents from the antimetabolite drug class, treating various malignancies, including leukemia, non-Hodgkin’s lymphoma, and cancers of the breast, prostate, lung, head and neck, and gastrointestinal tract [23–28]. However, both drugs have been shown to be neurotoxic [29–31]. Emerging evidence suggests that chemotherapeutic drugs, such as MTX and 5-FU, accelerate brain aging, with CRCI sharing pathological similarities with early-onset frailty and neurodegenerative diseases such as Alzheimer’s disease (AD)[32–34]. Given these parallels, interventions demonstrating neuroprotective effects in neurodegenerative conditions could potentially mitigate CRCI. One such intervention is hyperbaric oxygen (HBO) therapy.

HBO is currently an FDA-approved treatment for 13 medical conditions, including decompression sickness, air embolism, chronic wounds, and infections [35]. Conventional HBO therapy involves the administration of 100% oxygen at pressures 2.0-3.0 ATA for 90-120 minutes [36]. The therapeutic effects of HBO stem from increased hydrostatic pressure and high partial pressure of inspired oxygen. According to Boyle’s law, increased pressure reduces air bubble size, aiding in their dissolution and exhalation, benefitting conditions like air embolism [37]. Additionally, Henry’s law describes how higher partial pressure enhances oxygen dissolution in blood, thereby improving oxygen delivery to hypoxic tissues while enabling HBO’s secondary antimicrobial, anti-inflammatory, and angiogenic effects [38].

Pre-clinical and clinical studies demonstrated that HBO exposure improves cancer survival rates and boosts chemotherapy’s anti-tumor efficacy by overcoming tumor hypoxia, improving tumor perfusion, and increasing malignant cell sensitivity to destruction—all while maintaining safety in combination with cancer treatments [39–42]. Indeed, many chemotherapy drugs work by inducing oxidative stress in the tumors resulting in cell death, HBO enhances the killing of tumor cells by producing reactive oxygen species, contributing to the synergistic effects of HBO as an adjuvant cancer treatment [43–46]. Furthermore, HBO has shown neuroprotective effects in conditions such as Alzheimer’s disease, Parkinson’s disease, spinal cord injury, traumatic brain injury (TBI), and ischemic stroke [47, 48]. In neurodegenerative diseases, HBO is thought to prevent cognitive impairment and hippocampal pathology by counteracting cerebral hypoxia, improving microcirculation and neurogenesis, augmenting cholinergic pathways, reducing oxidative damage and neuroinflammation, exerting anti-apoptotic effects, and modulating epigenetic factors involved in cellular senescence [49]. Given its neuroprotective properties and its potential as an adjunct cancer therapy, HBO represents a promising candidate for CRCI intervention.

CRCI affects a substantial number of cancer survivors, yet its prevalence and severity across sexes remain unclear. Large-scale analyses of over 23,000 cancer patients (1980-2019) indicate that women have a 34% increased risk of experiencing severe treatment-related adverse events compared to men, with a 36% increased risk specifically for chemotherapy-induced toxicity [50]. Fatigue, a common chemotherapy side effect affecting up to 80% of individuals undergoing treatment [51], appears more prevalent in female patients [52, 53]. Persistent fatigue can impair attention, concentration, motivation, and functional abilities [54]. Additionally, geriatric female cancer patients scored lower on the Mini Mental State Examination (MMSE) compared to male patients [55]. While the precise sex-related prevalence of CRCI remains unclear, existing evidence highlights greater susceptibility to chemotherapy-induced toxicity in females [56]. Despite this, most preclinical studies on CRCI interventions have predominantly used male animals, limiting our understanding of treatment efficacy in females [57]. Given that chemotherapy is widely used across various cancers, including breast cancer, which disproportionately affects women, it is crucial to expand preclinical research to include both sexes to ensure comprehensive and effective treatment strategies.

This study explored the potential therapeutic effects of HBO against CRCI using a mouse model of MTX- and 5-FU-induced cognitive impairment. Additionally, we assessed whether the potential therapeutic effects of HBO vary depending on chemotherapy dosage and whether effects are observed in both sexes.

## 2. Methods

### 2.1 Ethics Statement

The NIH guidelines for the Care and Use of Laboratory Animals were followed in this study. Protocols containing procedures and handling of the mice were approved by the Institutional Animal Care and Use Committee at the University of North Texas Health Science Center (UNTHSC) (protocol #IACUC-2022-0030 approved on November 17, 2022).

### 2.2 Animals

Male and female C57BL/6J (Strain Number: #000664) mice were obtained from The Jackson Laboratory (Bar Harbor, ME) at 15 weeks of age and maintained at the UNTHSC Vivarium for 1 week of acclimation. The mice were subcutaneously inserted with radio-frequency identification chip (2-mm x 13-mm) for individual identification. All cages were maintained at 23 ± 1 °C, under a 12-h light/dark cycles starting at 0700, and given ad libitum access to food (ProLab® RMH 1800, LabDiet, #5LL2) and water. No environmental enrichment was provided in the cages as it could alter behavior and bias the outcomes of the study [58–60]. Animals were group housed (2-5 mice/cage based on sex and treatment) to provide social enrichment/interaction.

### 2.3 Treatments

#### Chemotherapy cocktails

Methotrexate hydrate (MTX, Millipore Sigma, CAS number 133073-73-1) and 5-fluorouracil (5-FU, MP Biomedicals, CAS number 51-21-8) were dissolved in 0.9% NaCl, adjusted to pH 7.0-7.4, and kept at 4°C.

#### Chemotherapy treatments (Weeks 1-3)

mice were randomly assigned to one of the experimental groups based (with each group having similar starting mean body weights within each sex). At 16 weeks of age, mice were randomly assigned to receive intraperitoneal (I.P.) injection of either 0.9% NaCl (Saline), or therapeutic cocktails of either low-dose (LD) (MTX 37.5 mg/kg and 5-FU 50 mg/kg) or high-dose (HD) (MTX 70 mg/kg and 5-FU 100 mg/kg) once a week for three weeks.

#### Hyperbaric oxygen therapy (Weeks 1-3)

Half of the mice were randomly assigned to HBO exposure, receiving one session per day (5 days/week, Monday to Friday) for three weeks. Mice, in their home cage, were placed inside a hyperbaric chamber and exposed to pure oxygen. The oxygen compression was gradually increased over 15 minutes, maintained at 2.4 ATA for 90 minutes, and gradually decompressed over 15 minutes. Mice not receiving HBO exposure were also brought into the room to minimize travel/room/ environment effects.

For each sex, there are six treatment groups: Saline, Saline + HBO, LD, LD + HBO, HD, and HD + HBO. Experimental design and timeline are summarized in schematic Figure 1.

**Figure 1.**
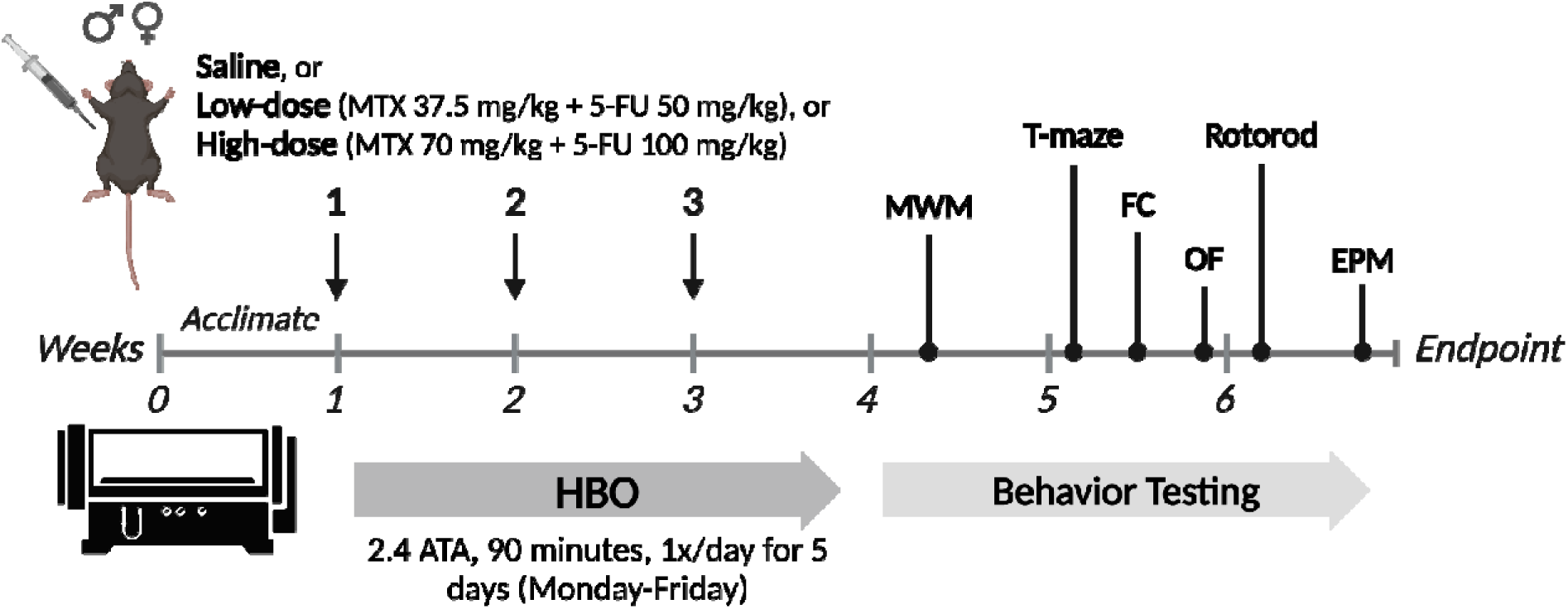
Schematic diagram of experimental procedure. Mice were given intraperitoneal injections of either saline or methotrexate (MTX) and 5-fluorouracil (5-FU) at low-dose (LD) and high-dose (HD) to induce chemotherapy-related cognitive impairment (CRCI), once a week (weeks 1-3). Concurrently, daily hyperbaric oxygen (HBO) therapy was given to subsets of mice 5 days a week (weeks 1-3). Behavior testing was conducted (weeks 4-6): Morris water maze (MWM), active avoidance (T-maze), fear conditioning (FC), open field (OF), rotorod, and elevated plus maze (EPM). Created in BioRender. Trinh, K. (2025). https://BioRender.com/o45d574.

### 2.4 General health assessment

Body weights, clinical observation, grip strength, wire suspension, and food intake were recorded weekly throughout the study.

#### 2.4.1 Clinical observations

A 13-item examination was adapted from a previously established 31-item frailty index to assess clinical signs of treatment-induced deterioration [61]. Specifically, the integument (alopecia, loss of fur color, dermatitis, loss of whiskers, coat condition), the physical/musculoskeletal system (tumors, tremor, body condition), the auditory system (hearing loss), the ocular system (vision loss), the digestive system (diarrhea), and discomfort (piloerection, mouse grimace scale [62]) were evaluated on the scale of 0 (absent), 0.5 (mild), and 1 (severe). Each mouse was evaluated once weekly.

#### 2.4.2 Grip strength

Grip strength (GS) is a non-invasive way to assess the neuromuscular function of a mouse’s forelimbs [63, 64] as well as a clinically relevant index to assess treatment-induced frailty [65]. GS was measured using a commercial automatic grip strength meter (Bioseb, Vitrolles, France). Each mouse was gently lifted by the tail, then lowered until they grasped the metal grid with both forepaws. The mouse would then be gently and uniformly pulled horizontally relative to the force meter until it lost its grip and released the grid. The maximum resistance force were recorded (grams of weight), averaged across 3 trials, and normalized against the mouse’s body weight.

#### 2.4.3 Wire suspension

Muscle weakness, another clinical frailty measure, can be assessed with forelimb wire suspension [66]. Using a horizontal wire suspended 40-cm above a padded surface, the mouse was held gently by tail and allowed to grip onto the wire using two front paws. The latency to tread (grabbing the wire with either hind legs) and fall (maximum 30 seconds) from the wire were recorded. Each mouse was given one trial weekly.

## 3. Behavioral assessment

One week following the last injection and 3 days following the last HBO session, mice underwent behavioral evaluation of cognitive, affective and motor functions for three weeks (Weeks 4-6) in the following order:

### 3.1 Spatial learning and memory

Hippocampal-dependent spatial learning and reference memory were assessed using Morris water maze (MWM) [67]. A circular tank (120 cm in diameter, 50 cm deep) was filled with water opacified with non-toxic white paint and maintained at 24 ± 1°C. The Morris water maze task consisted of three components: (i) Pre-training, (ii) Spatial discrimination, and (iii) Probe trials.

i. During pre-training, mice were placed at one end of a straight alley and trained to swim and stand on the hidden platform located at the opposite end of the alley. All visual cues were removed by placing a black curtain over the tank. Mice were given 5 trials per session, 1 daily session for 2 days.
ii. During spatial discrimination, the curtain was removed and mice used distal visual cues in the testing room to locate the platform (10 cm x 10 cm) hidden 1.5 cm below the surface of water. Each session consisted of 5 trials, each starting from one of four possible starting locations, in which the mouse swam until it located the platform, or the maximum duration (90 s) had been reached. If the mouse did not locate the platform after 90 s, a test examiner would guide the mouse to the platform where it remained for 10 s before being placed in the holding cage for a 90-s intertrial interval. Mice were given 5 trials per session, 1 daily session for 4 consecutive days.
iii. Probe trials, given on the last trial of sessions 2 and 4 of spatial discrimination, involved removing the platform and allowing mice to explore for 30 s. At the end of which, the platform was brought back and mice were given time to find it.

Performance of mice, consisting of pathlength, latency, and speed, were tracked by a commercial tracking system (ANY-maze, Stoelting Co, IL, USA).

To study how treatments may influence an animal’s ability to solve a spatial task, search strategies during MWM were analyzed with Pathfinder (Jason Snyder Lab, Vancouver, Canada) [68]. Path maps from ANY-maze software were imported into Pathfinder and each MWM trial was assigned to one of the eight possible search strategies based on previously described criteria (Ideal path error, heading error, time in target zone, distance to goal, etc.) [69]. As described previously [70], search strategies are categorized into non-spatial and spatial. Non-spatial search category is less dependent on hippocampal functions, including ‘thigmotaxis’, ‘random search’, ‘scanning’, and ‘chaining’. On the contrary, spatial search category relies more on hippocampal functions, including ‘indirect search’, ‘focal search’, ‘directed search’, and ‘direct path’.

Schematic examples of the search strategies are demonstrated in schematic Figure 4. Incidence of search strategies was displayed as percentage of total trials to account for group size differences.

### 3.3 Learning and cognitive flexibility (active avoidance)

The active avoidance test evaluates the acquisition of associative learning and procedural memory, and spatial working memory. The test consists of a T-shaped maze featuring a start box (stem) and two lateral goal arms resting on a stainless-steel floor wired for scrambled shock (0.69 mA) to the feet. During first session (Acquisition), an initial preference trial is given wherein a shock is initiated in the start box and then terminated upon mouse’s entry to either of the lateral goal arm. The correct goal arm is then determined the arm opposite to the initially chosen arm. On each trial thereafter, shock will be initiated 5 seconds after the opening of the start box’s door or immediately upon entry into the incorrect arm and will be terminated after entry into the correct goal arm. Once inside the correct goal arm, mice stay in there for 10 seconds before returning to a temporary holding cage for 1-min ITI. Components of this test are scored independently, Avoidance: entering either of lateral arms within 5 seconds; Correct Discrimination: entering the correct goal arm without error, regardless of time requirement; Correct Avoidance: entering the correct goal arm within 5 seconds. The session continues until the mouse has reached criterion by making Correct avoidance on four out of five consecutive trials, with the last two trials are Correct Avoidance. One hour after the completion of Acquisition session is the Reversal session wherein the correct goal arm is then switched to the arm opposite to the correct arm in Acquisition session. Mice are given a maximum of 60 seconds per trials and a maximum of 25 trials per session to reach criterion. Number of trials to reach the Avoidance criterion is analyzed to evaluate the cognitive performance of the animals.

### 3.4 Fear conditioning

Fear conditioning (FC) was used to assess contextual and cued memory [71]. This form of associative learning requires the pairing of a neutral stimulus (conditioned stimulus, CS) with a foot shock (unconditioned stimulus, US). As mice learn to associate between the environment, the CS, and the US, mice produce a behavioral response such as freezing when encountering the environment or the CS. The FC apparatus consisted of plexiglass walls (19 x 18 x 24 cm) resting on a stainless-steel floor wired for scrambled shock to the feet. The apparatus was situated within a dimly lit (35 lux), sound-attenuated chamber equipped with a fan for ventilation and background noise. A camera was mounted in the ceiling of the chamber to track the freezing behavior (ANY-maze, Stoelting, Chicago, IL, USA). FC consisted of 4 sessions. Session 1 involved placing the mouse in an unfamiliar environment (Conditioning Context, CC) consisting of a grid floor, black-and-white stripe walls, and vinegar smell (distilled white vinegar 5% acidity) for a 5-min duration, within which 2 pairings of loud sound lasting 15 seconds (Conditioned Stimulus (CS); 2,000 Hz) and a foot shock (Unconditioned Stimulus (US); 0.69 mA) occurring during the last 2 seconds of CS were presented and separated by 2 min. 24 hours later was session 2, in which the mouse was returned to the environment (CC) same as that of session, but without CS or US, for a 5-min duration. One hour later was session 3, the mouse was presented with a novel environment (Novel Context, NC) consisting of a smooth floor, grey walls, and alcohol smell (70% EtOH) for a 3-min duration. Immediately after session 3 was session 4, while the mouse was still in NC, the loud sound CS was presented for a 3-min duration. Percentage of time freezing in each session was recorded by ANY-maze (Stoelting) for analysis. Each mouse was tested once on the fear conditioning test.

### 3.5 Spontaneous locomotor activity

Spontaneous locomotion and anxiety-related behaviors were assessed via open field test using a Digiscan apparatus (Omnitech Electronics, model RXYZCM). The apparatus consisted of a clear acrylic cage (40.5 x 40.5 x 30.5 cm) enclosed by a metal frame lined with photocells. The apparatus was situated within a chamber that provided dim light (23 lux) and background noise (80 dB). For a 16-min duration, each mouse was let freely explored inside the apparatus, movements were tracked by photocells and processed by software program (Fusion v5.5 Superflex Edition, Omnitech Electronics, Columbus, OH, USA).

### 3.6 Coordinated running

Motor learning and coordination were assessed using an accelerating rotorod test (AccuRotor, Omnitech Electronics, Columbus, OH, USA) consisting of a 3.2-cm diameter nylon cylinder mounted horizontally at 35.5 cm above a well-padded surface. In each trial, each mouse was placed on the cylinder, which began to rotate at increasing speed (accelerating from 0-750 RPM in 150 seconds) until the mouse fell to the padded surface below, each mouse’s latency to fall were recorded. Each session consisted of 4 trials with 10-min ITI. Mice were given 2 sessions per day, separated by 2-3 hours, until the running performance has reached a criterion (three consecutive sessions by which the average latency to fall did not differ by more than 15%) at the end of session 7 or thereafter.

### 3.7 Anxiety-like behavior

Elevated plus maze (EPM) could assess anxiety-related behaviors [72]. The maze was made of plastic, consisted of four arms (two arms open to the room and 2 arms enclosed with plexiglass walls such that the floor was not visible) 30-cm long and 5-cm wide creating the plus sign “+” shape, with the same type of arms facing each other. The maze was elevated 45-cm above the floor and directly under a 60-watt spotlight in a dark room. For a 5-min duration, each mouse freely explored the maze. A camera mounted above the maze tracked mouse movements, and time spent in each arm was recorded and processed using a software program (ANY-maze, Stoelting, Chicago, IL, USA).

## 4. Statistical Analysis

Statistical analyses were performed with SYSTAT 13 (Version 13.1) (SYSTAT Software, Inc. 2009, San Jose, CA). Data are presented as mean ± standard error of mean (S.E.M). Body weights, clinical observations, grip strength, wire suspension, food intake, and each search strategy results were analyzed using repeated measure two-way analyses of variance (ANOVA) with between group factors of chemotherapy (Chemo), HBO intervention (Int), and week or session as repeated measure. Behavioral performance was analyzed using two-way ANOVA with chemotherapy and HBO intervention as factors. Significant main effects or interaction were followed by post-hoc pairwise comparison of means using Fisher’s Least Significant Difference to assess group differences. The α level was set at 0.05 for all analyses.

## 5. Results

### 5.1. General health assessment

#### 5.1.1. Body weights and food intake

In males (Fig. 2A), body weights from HD, but not LD, decreased over the first 4 weeks but returned to initial weight for both +/- HBO groups, supported by a main effect (F_CHEMO_ (2, 86) = 3.194; *p* = 0.046) and interactions (F_WEEK*CHEMO_ (10, 430) = 9.005; *p* < 0.001; F_WEEK*INT_ (5, 430) = 9.005; *p* = 0.048). Similarly, in females (Fig. 2B), HD, but not LD, decreased body weights over the first 4 weeks but recovered by the end of study, whereas co-treatment with HBO mitigated this transient weight loss, supported by interaction (F_WEEK*CHEMO*INT_ (10, 390) = 5.68; *p* < 0.001).

**Figure 2.**
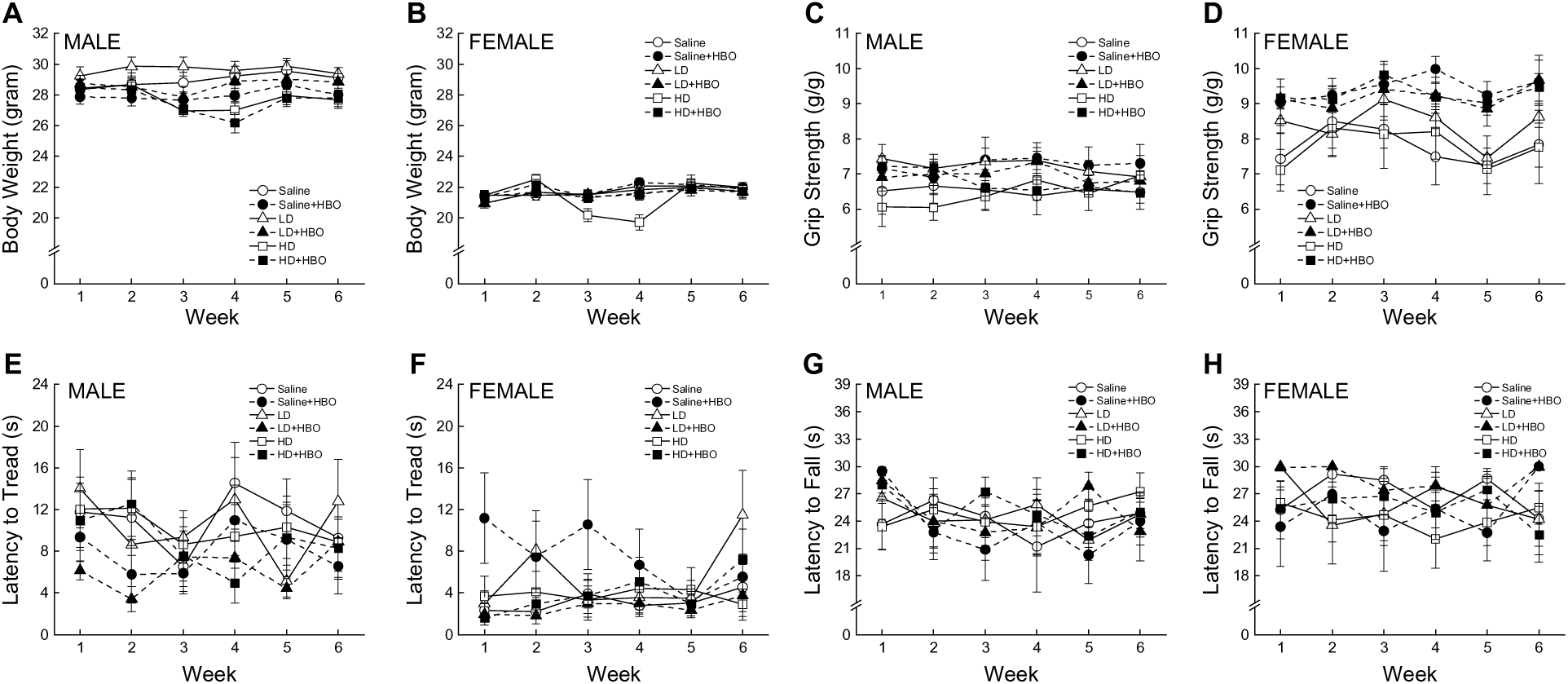
Effects of chemotherapy and HBO on general health. Body weights of males (**A**) (n=14-17) and females (**B**) (n=13-15). Grip strength normalized to body weights in males (**C**) (n=10-17) and females (**D**) (n=8-15). Wire suspension: latency to tread (**E**, **F**) and latency to fall (**G**, **H**) in males (n=10-17) and females (n=8-15). Data are presented as mean ± S.E.M.

In males (Supplementary Figure 1A), all groups’ average weekly food intake decreased gradually throughout the 6 weeks, supported by a main effect (F_WEEK_ (5, 90) = 10.157; *p* < 0.001) and HD chemotherapy decreased consumption compared to other groups during treatment (weeks 1-3), as supported by main effect (F_CHEMO_ (2, 18) = 5.87; *p* = 0.011) and interaction (F_WEEK*CHEMO_ (10, 90) = 2.916; *p* = 0.003). In females (Supplementary Figure 1B), average weekly food intake also decreased gradually (F_WEEK_ (5, 85) = 19.493; *p* < 0.001), but neither chemotherapy or HBO intervention affected food intake (all Fs < 0.860, all *ps* > 0.570).

#### 5.1.2. Clinical observations

In males (Supplementary Figure 2A), alopecia score from LD, but not HD, increased over weeks 2-3 but returned to initial level by the end of the study, supported by a main effect (F_CHEMO_ (2, 64) = 6.972; *p* = 0.002) and an interaction (F_WEEK*CHEMO_ (10, 320) = 3.763; *p* < 0.001). HBO seemed to prevent LD from alopecia recovery during the last 3 weeks, as supported by an interaction (F_WEEK*INT_ (5, 320) = 2.257; *p* = 0.049). Similarly (Supplementary Figure 2C), LD had worse coat condition scores throughout the study, supported by an interaction (F_WEEK*CHEMO_ (10, 320) = 3.159; *p* = 0.001), and HBO reversed this effect as supported by main effect (F_INT_ (1, 64) = 5.236; *p* = 0.025) and interaction (F_WEEK*CHEMO*INT_ (10, 320) = 1.978; *p* = 0.035). Visually, dermatitis score did not seem to be impacted by chemotherapy of HBO (Supplementary Figure 2E- lack of variation/effect in most groups did not allow for statistical comparisons). In females, neither chemotherapy or HBO induced alopecia, dermatitis, or impaired coat condition (Supplementary Figure 2B, D, F- lack of variation/effect in most groups did not allow for statistical comparisons).

#### 5.1.3. Grip strength

In males (Fig. 2C), there were no significant effects of chemotherapy doses or HBO on grip strength over the course of the study (all Fs < 1.600, all *ps* > 0.130). In females (Fig. 2D), HBO groups seemed to have higher grip strength from the start and throughout the study, as supported by a main effect (F_INT_ (1, 54) = 10.543; *p* = 0.002).

#### 5.1.4. Wire suspension

In males (Figs. 2E and 2G), there were no significant effects of chemotherapy doses or HBO on wire suspension performance across the study (all Fs < 3.760, all *ps* > 0.055). In females (Figs. 2F and 2H), saline + HBO group had higher latency to tread compared to the other group over the first 4 weeks of the study. This observation was supported by a significant interaction (F_WEEK*CHEMO*INT_ (10, 270) = 2.034; *p* = 0.03).

### 5.2 Spatial learning and memory

#### 5.2.1. Initial Learning (Sessions 1-2)

In males (Figs. 3A,3C, 3D), there was no significant effect of chemotherapy or HBO on path length taken to reach the platform or speed, supported by a lack of main effect or interaction (all Fs < 3.74, all *ps* > 0.05). However, HD groups had higher latencies, especially HD + HBO. This observation was supported by a main effect (F_CHEMO_ (2, 85) = 3.871; *p* = 0.025). In females (Figs. 3B, 3D, 3F), there was no significant effect of chemotherapy or HBO on path length (all Fs < 2.57, all *ps* > 0.05). However, HD groups had higher latencies and lower speeds. These observations were supported by main effects (Latency: F_CHEMO_ (2, 78) = 10.093; *p* < 0.001; Speed: F_CHEMO_ (2, 78) = 9.29; *p* < 0.001).

**Figure 3.**
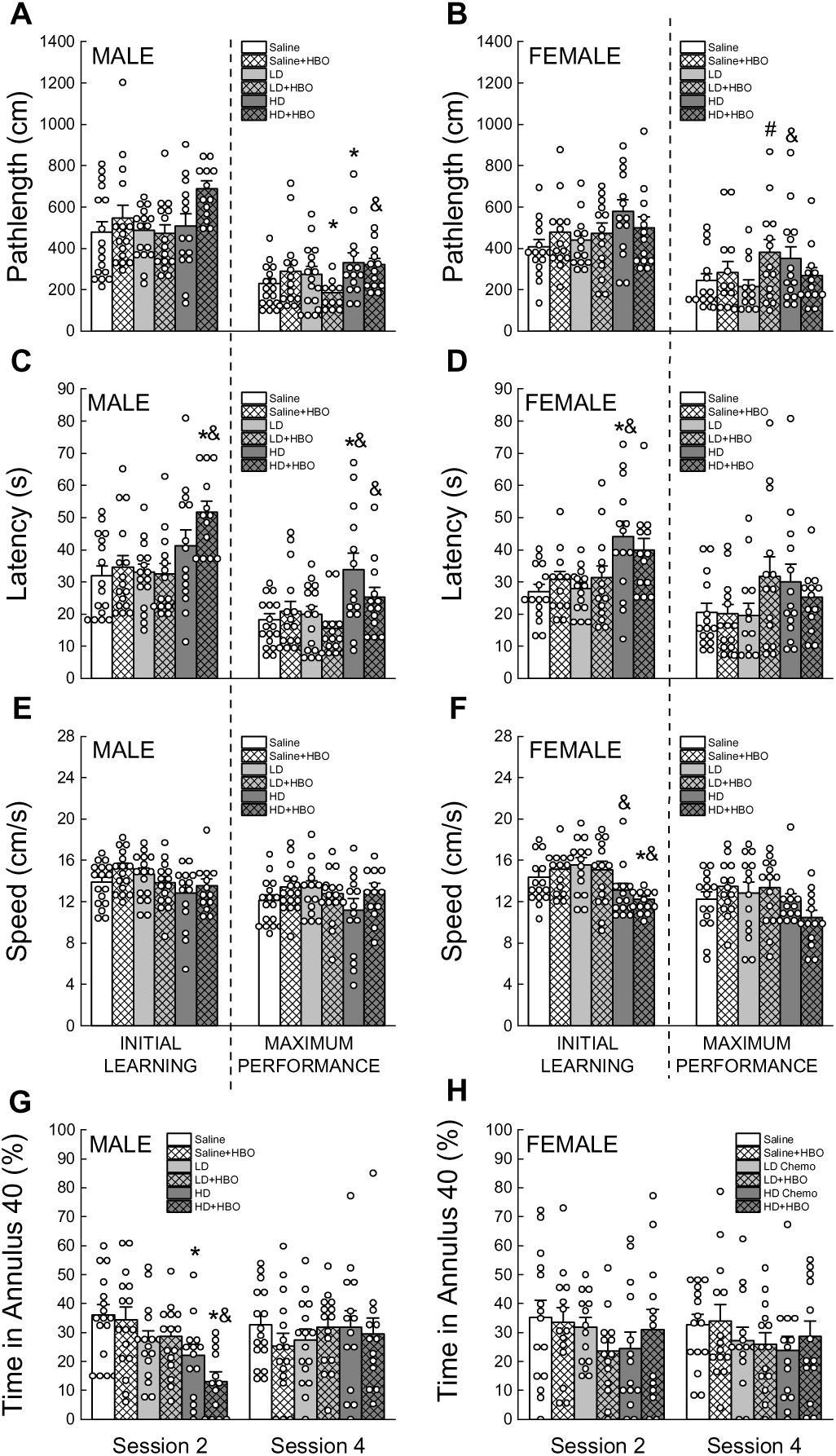
Effects of chemotherapy and HBO on spatial learning and memory. Performance was assessed as pathlength (**A**, **B**), latency (**C**, **D**), and swimming speed (**E**, **F**) during initial learning (average sessions 1&2) and maximum performance (average sessions 3&4) in male (left panels) and female (right panels) mice. Probe performance was measured as time spent in Annulus 40 area surrounding where hidden platform was placed (**G**, **H**). Data are presented as mean ± S.E.M (Male: n=13-17; Female: n=13-15). * p<0.05 versus intervention-matched Saline; & p<0.05 versus intervention-matched LD; # p<0.05 versus chemotherapy-matched non-HBO.

**Figure 4.**
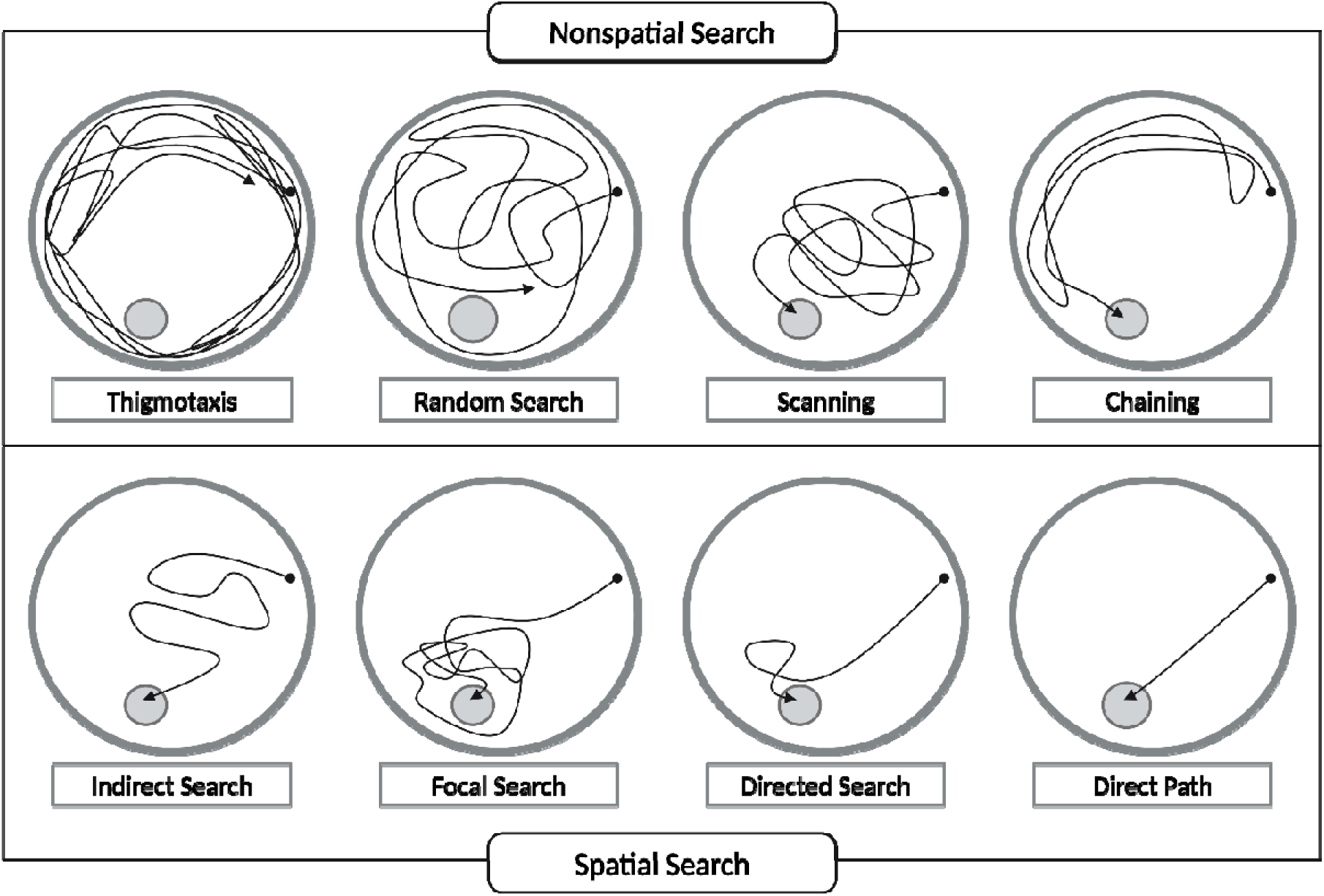
Representative swim paths of search strategies (Morris water maze). Eight search strategies are categorized as <u>nonspatial</u> (thigmotaxis, random search, scanning, and chaining) and <u>spatial</u> (indirect search, focal search, directed search, and direct path). Swim paths illustrate how mice, from starting point (black dot), use spatial cues to locate the submerged platform (grey circle). Created in BioRender. Trinh, K. (2025) https://BioRender.com/z92g102.

#### 5.2.2. Maximum Performance (Sessions 3-4)

In males (Figs. 3A,3C, 3D), path length and latency were higher in HD groups, supported by main effects (Path Length: F_CHEMO_ (2, 85) = 3.871; *p* = 0.025; Latency: F_CHEMO_ (2, 85) = 8.26; *p* = 0.001). Speed was not affected by either chemotherapy or HBO (all Fs < 2.3, all *p*s > 0.13). In females (Figs. 3B, 3D, 3F), LD +HBO and HD alone had higher path length than the other groups, which was supported by an interaction (F_CHEMO*INT_ (2, 78) = 3.271; *p* = 0.043). Neither speed nor latency were affected by chemotherapy or HBO (all Fs < 2.90, all *p*s < 0.07).

#### 5.2.3. Probe trials

In males (Fig. 3G), HD groups spent less time in the annulus 40 during session 2, which was supported by a main effect (F_CHEMO_ (2, 85) = 11.616; *p* < 0.001). There was no effect of chemotherapy or HBO during session 4 (all Fs < 0.27, all *p*s > 0.384). In females (Fig. 3H), there was no significant effect of chemotherapy or HBO on either session (all Fs < 0.5, all *p*s > 0.230)

#### 5.2.4. Search strategies

All treatment groups progressed towards utilizing less non-spatial strategies (‘thigmotaxis’, ‘random search’, ‘scanning’, and ‘chaining’) and more spatial strategies (‘indirect search’, ‘focal search’, ‘directed search’, and ‘direct path’) over 4 sessions (Fig. 5), as supported by a main effect (Male & Female: All F(s)_SESSION_ > 3.370; *ps* < 0.001).

**Figure 5:**
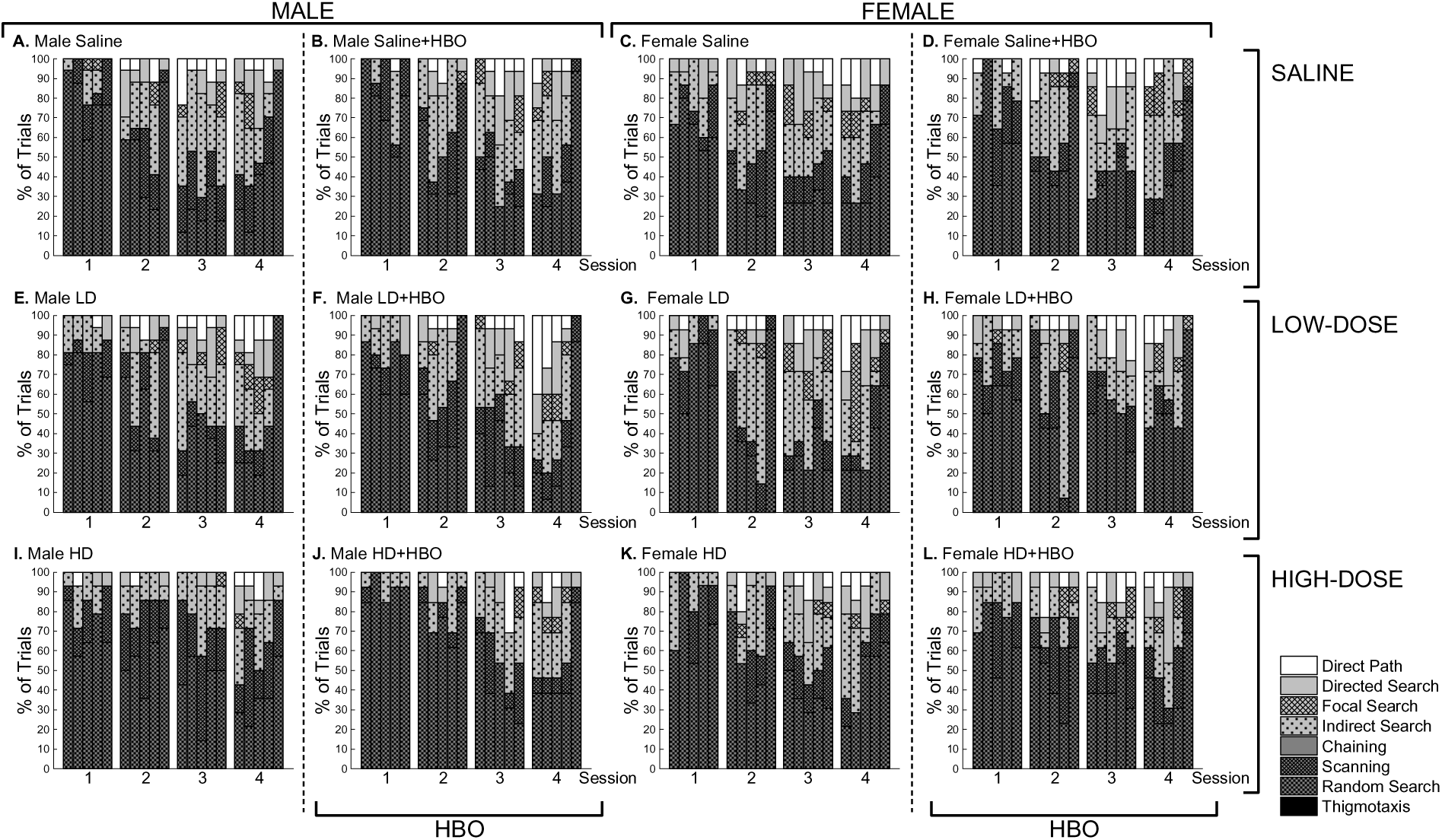
Effects of chemotherapy and HBO on Morris water maze navigation. Distribution of search strategies used by saline (**A, C**), low-dose (**E, G**), high-dose (**I, K**), and co-treatment of HBO with saline (**B, D**), low-dose (**F, H**), and high-dose (**J, L**) in male (left panels) and female (right panels) mice. Data are presented as the percentage of search strategies utilized in each trial of 4 sessions (Male: n=13-17; Female: n=13-15).

In males (Fig. 5 & Supp. Table 1, left panels), chemotherapy, especially HD, groups relied more on non-spatial ‘random search’ strategies than Saline groups especially during sessions 2 and 3 (Supp. Table 1. **Session 2**, Sal: 42.6% vs. LD: 48.4% vs. HD: 51.8%; **Session 3**, Sal: 21.2% vs. LD: 35% vs. HD: 42.9%), supported by a main effect (F_CHEMO_ (2, 85) = 5.537; *p* = 0.005). HBO reduced the use of ‘random search’ in the LD groups during sessions 2-4 (Supp. Table 1. **Session 2**, LD: 48.4% vs. LD+HBO: 38.3%; **Session 3**, LD: 35.0% vs. LD+HBO: 26.7%; **Session 4**, LD: 28.1% vs. LD+HBO: 18.3%), but increased its use in the HD group (Supp. Table 1. **Session 1**, HD: 70% vs. HD+HBO: 90.8%; **Session 2**, HD: 51.8% vs. HD+HBO: 73.1%), however the interaction did not reach significance (F_CHEMO*INT_ (2, 85) = 3.08; *p* = 0.051). ‘Scanning’, a non-spatial strategy at higher level than ‘random search’, was not affected by chemotherapy (F_CHEMO_ (2, 85) = 1.076; *p* = 0.345). However, HBO exposure influenced ‘scanning’ use dependent on chemotherapy doses. While HBO had minimal effect on LD, HBO significantly reduced utilization of ‘scanning’ in HD groups throughout 4 sessions (Supp. Table 1. **Session 1**, HD: 14.3% vs. HD+HBO: 1.5%; **Session 2**, HD: 28.6% vs. HD+HBO: 5.8; **Session 3**, HD: 30.0% vs. HD+HBO: 15.4%; **Session 4**, HD: 26.8% vs. HD+HBO: 9.6%), supported by a main effect (F_INT_ (1, 85) = 8.165; *p* = 0.005) and interaction (F_CHEMO*INT_ (2, 85) = 8.733; *p* < 0.001). The ‘direct path’ strategy, which represents the highest level of hippocampal-dependent spatial search, was is used more by the LD group only during session 4, and it was further increased in LD+HBO groups (Supp. Table 1. **Session 4**, Sal: 5.9% vs LD: 14.1% vs. HD: 7.1%; LD vs. LD+HBO: 21.7%). This was supported by a main effect (F_CHEMO_ (2, 85) = 3.932; *p* = 0.023) and interaction (F_SESSION*CHEMO_ (6, 255) = 3.039; *p* = 0.007). No other search strategies were affected by treatments in males (all Fs < 2.627, all *ps* > 0.077).

In females (Fig. 5 & Supp. Table 1, right panels), all treatment groups exhibited similar incidence of search strategies over 4 sessions, supported by lack of main effects (all Fs < 2.306, all *ps* > 0.105).

#### 5.2.5. Hippocampal-dependent spatial search

In males (Fig. 6A), HD significantly lowered the utilization of spatial search strategies throughout MWM sessions, supported by a main effect (Initial learning: F_CHEMO_ (2, 85) = 4.261; *p* = 0.017; Maximum performance: F_CHEMO_ (2, 85) = 4.449; *p* = 0.015). HBO did not mitigate this reduction in spatial navigation (all Fs < 1.035, all *p*s > 0.310). In females (Fig. 6B), there was a trend of HD groups using fewer spatial strategies during initial learning, however the main effect did not reach significance (F_CHEMO_ (2, 78) = 3.026; *p* = 0.054). There was no effect of HBO during initial learning, and there was no effect of chemotherapy nor HBO during maximum performance (all Fs < 1.154, all *p*s > 0.285).

**Figure 6.**
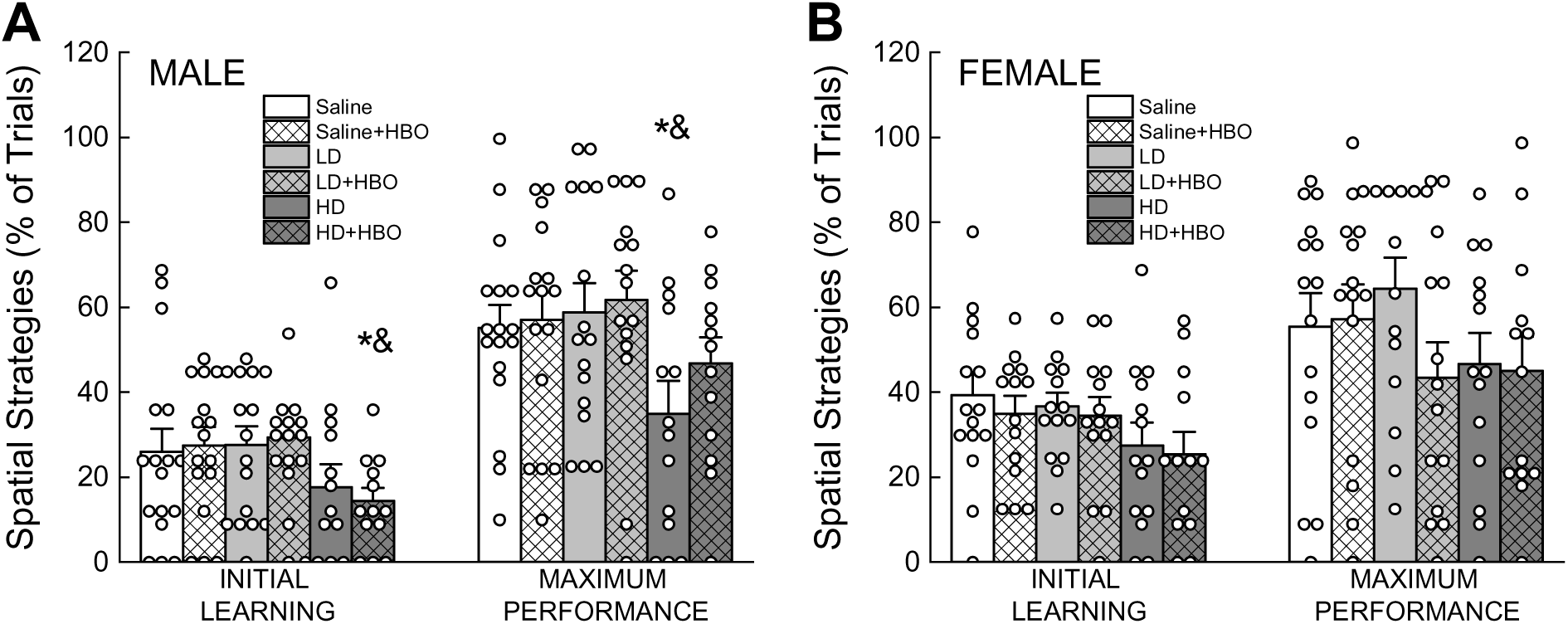
Effects of chemotherapy and HBO on spatial navigation (Morris water maze). Spatial search includes indirect search, focal search, directed search, and direct path strategies. Utilization of spatial search (percentage of total trials) during initial learning (average sessions 1&2) and maximum performance (average sessions 3&4, excluding probe trials) male (**A**) and female (**B**) mice. Data are presented mean ± S.E.M (Male: n=13-17; Female: n=13-15). * p<0.05 versus intervention-matched Saline; & p<0.05 versus intervention-matched LD.

### 5.4 Learning and cognitive flexibility

In males (Fig. 7A), LD mice took more trials to reach criterion during acquisition, which led to a significant main effect (F_CHEMO_ (2, 85) = 3.129; *p* = 0.049) (Fig. 4A). HBO significantly improved performance regardless of chemotherapy, which was supported by a main effect (F_INT_ (2, 85) = 3.129; *p* < 0.001). There was no effect of chemotherapy or HBO during reversal (all Fs < 1, all *p*s > 0.380). In females (Fig. 7B), chemotherapy-only groups, especially HD, took more trials to reach criterion than Saline, and co-treatment with HBO significantly reduced number of trials compared to LD- and HD-only groups. This observation during acquisition was supported by a significant interaction (F_CHEMO*INT_ (2, 78) = 4.199; *p* = 0.019). During reversal, HD+HBO took less trials to reach criterion than the HD group, supported by a significant interaction (F_CHEMO*INT_ (2, 78) = 3.339; *p* = 0.041).

**Figure 7.**
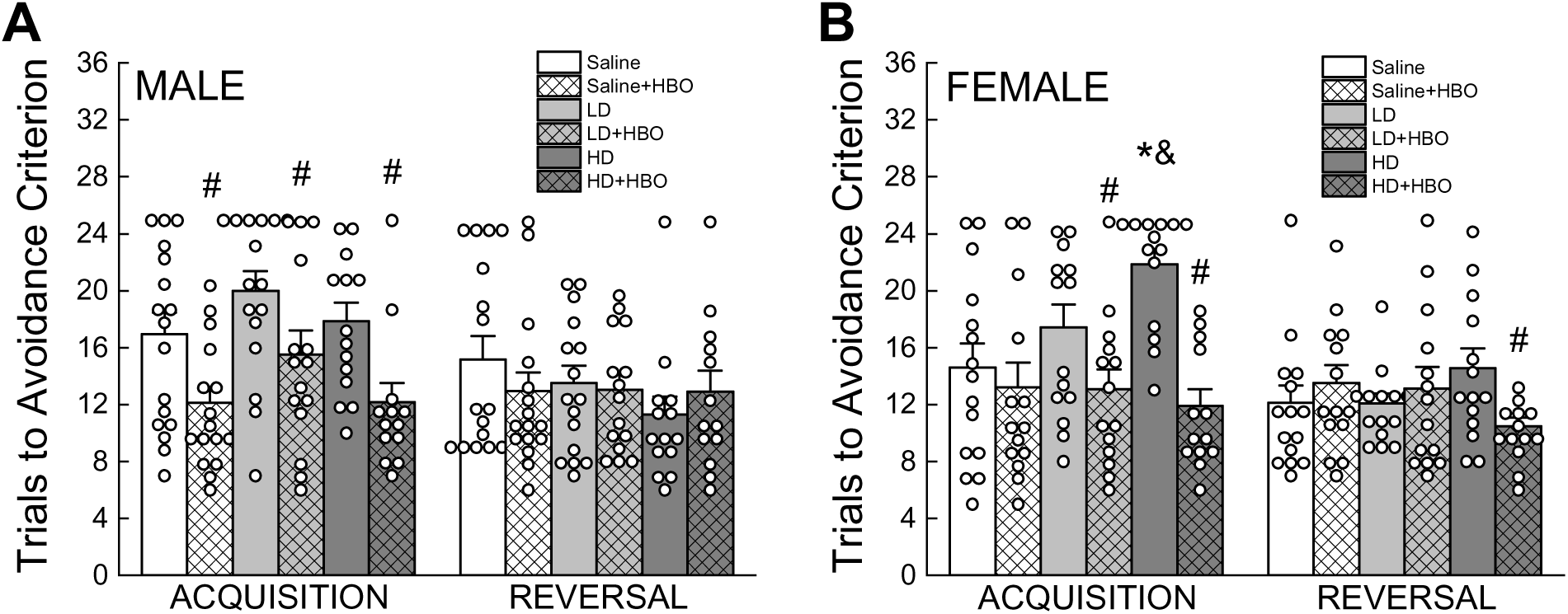
Effects of chemotherapy and HBO on avoidance learning (acquisition) and cognitive flexibility (reversal) (T-maze). Bar graphs depict number of trials to reach Avoidance criterion in male (**A**) and female (**B**) mice. Data are presented mean ± S.E.M (Male: n=13-17; Female: n=13-15). * p<0.05 versus intervention-matched Saline; & p<0.05 versus intervention-matched LD; # p<0.05 versus chemotherapy-matched non-HBO.

### 5.5 Fear conditioning

#### 5.5.1. Novel context (NC)

In males (Fig. 8A), HD spent more time freezing compared to saline, which was supported by a main effect (F_CHEMO_ (2, 85) = 3.654; *p* = 0.03). There was no effect of HBO on performance (F_INT_ (1, 85) = 0.262, *p* = 0.61). In females (Fig. 8B), there was no significant effect of chemotherapy or HBO on freezing behavior (all Fs <3.0, all *p*s > 0.085).

**Figure 8.**
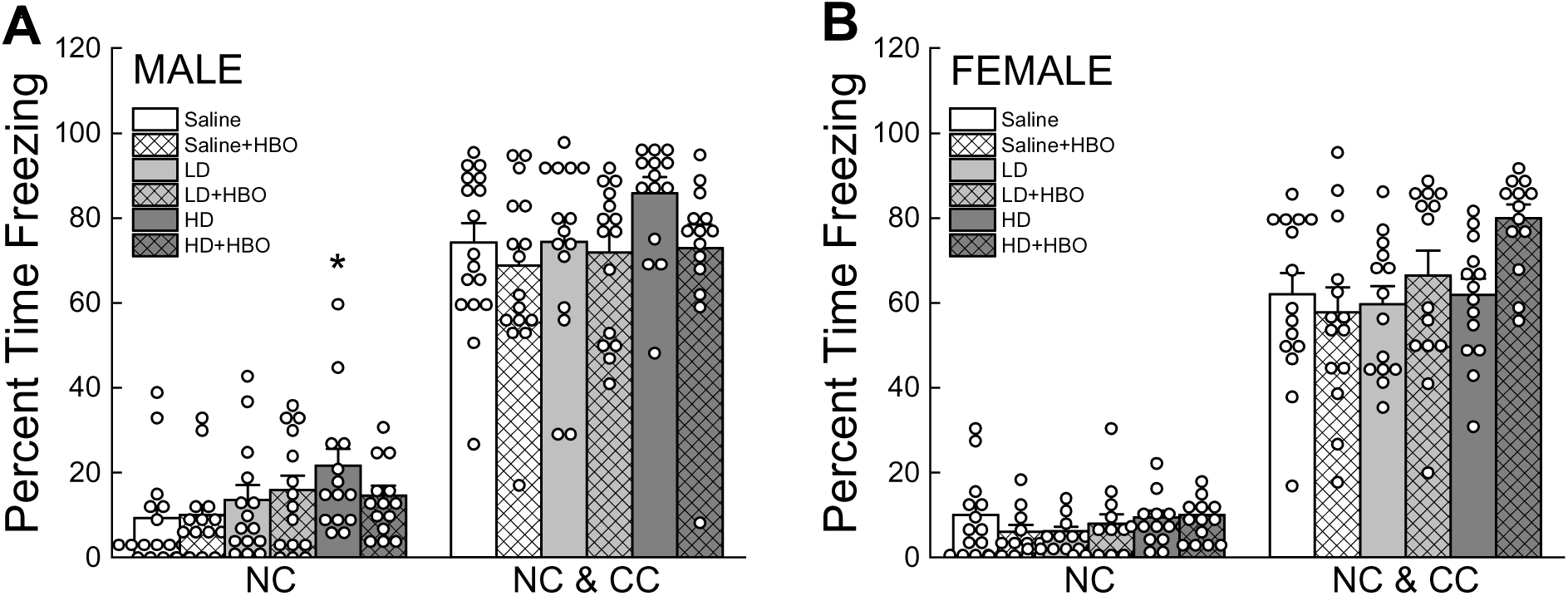
Effects of chemotherapy and HBO on contextual and cued memory (fear conditioning). Bar graphs showing prevalence of freezing behavior (percentage of time) in novel context (NC) (left) and when conditioning stimulus was added (NC & CS) (right) in male (**A**) and female (**B**) mice. Data are presented mean ± S.E.M (Male: n=14-17; Female: n=13-15). * p<0.05 versus intervention-matched Saline; & p<0.05 versus intervention-matched LD; # p<0.05 versus chemotherapy-matched non-HBO.

#### 5.5.2. Novel Context and Conditioned Stimulus (NC & CC)

In males (Fig. 8A), there was no significant effect of chemotherapy or HBO on freezing behavior (all Fs <1.5, all *p*s > 0.25). In females (Fig. 8B), there was no significant effect of chemotherapy or HBO on freezing behavior (all Fs <2.99, all *p*s > 0.070).

### 5.6 Spontaneous locomotor activity

In males (Figs. 9A, 9C, 9E), LD + HBO and both HD groups travelled less distance than the other groups, which was supported by a main effect (F_CHEMO_ (2, 63) = 7.587, *p* = 0.001). There was a trend for these groups to have less vertical activity but the main effect did not reach significance (F_CHEMO_ (2, 63) = 2.483, *p* = 0.092). LD and HD groups spent less time in the center, which was supported by a main effect (F_CHEMO_ (2, 63) = 3.593, *p* = 0.033). In females (Figs 9B, 9D, 9F), there was no effect of either chemotherapy or HBO intervention in distance travelled and in vertical activity (all Fs < 1.50, all *p*s > 0.240). While LD or HD alone had no effect on time spent in the center, LD/HD + HBO spent less time in that zone, which was supported by a main effect (F_INT_ (1, 54) = 4.883, *p* = 0.031) and an interaction (F_CHEMO*INT_ (2, 54) = 3.77, *p* = 0.029).

**Figure 9.**
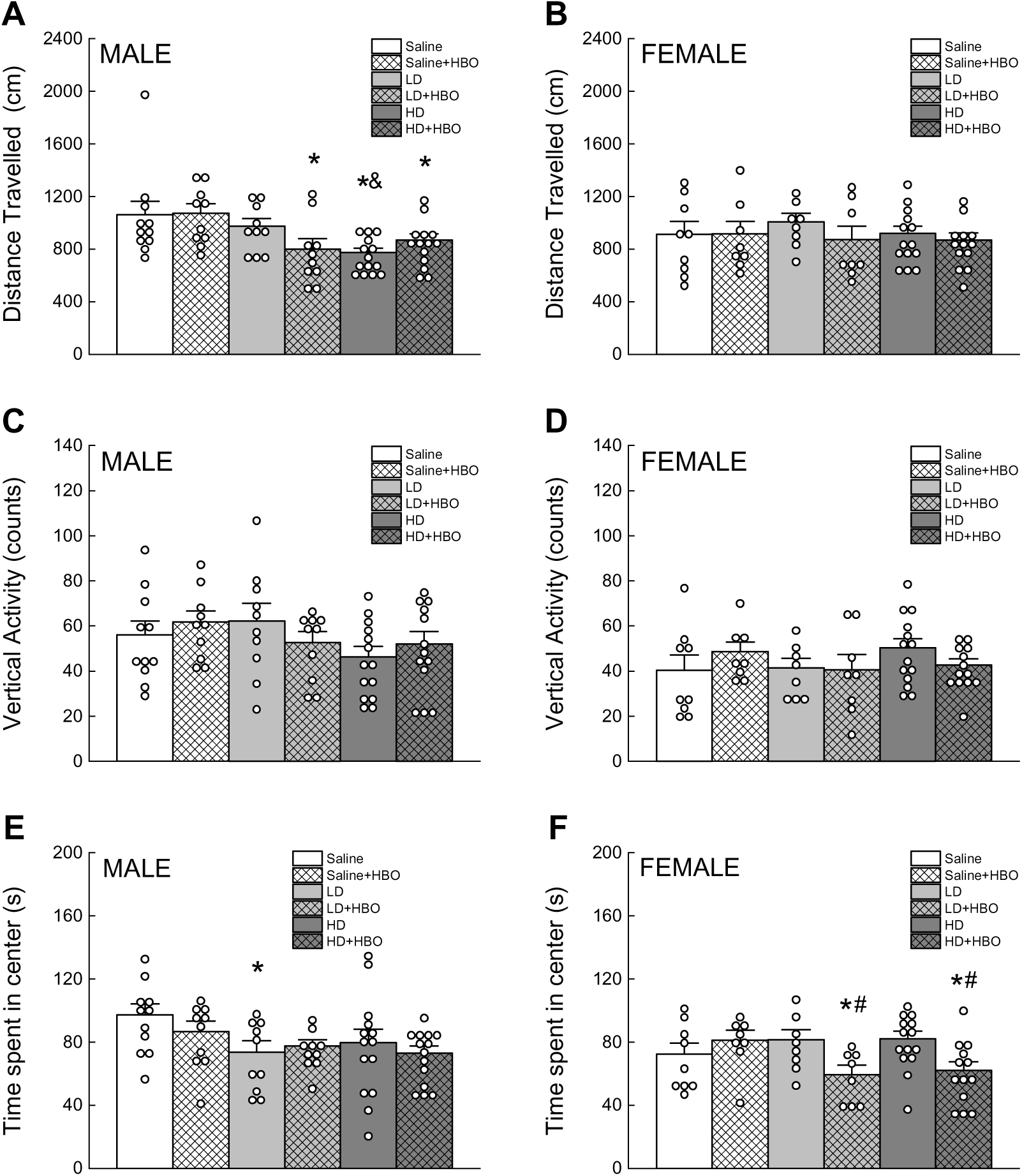
Effects of chemotherapy and HBO on spontaneous activity (open field). In a 16-minute test duration, the average travelled distance (**A**, **B**), vertical activity counts (**C**, **D**), and time spent in the center area (s) (**E**, **F**) was calculated for male (left panels) and female (right panels) mice. Data are presented mean ± S.E.M (Male: n=10-14; Female: n=8-14). * p<0.05 versus intervention-matched Saline; & p<0.05 versus intervention-matched LD; # p<0.05 versus chemotherapy-matched non-HBO.

### 5.7 Coordinated running

In males (Figs. 10A, 10C, 10E), there was no effect of either chemotherapy or HBO during initial learning (all Fs < 2.200, all *ps* > 0.100). During sessions 4-7, HBO seemed to lower latency in LD and increase it in HD, which was supported by a chemotherapy by intervention interaction (F_CHEMO*INT_ (2, 85) = 3.163, *p* = 0.047). The same observation was made for maximum performance, also supported by an interaction (F_CHEMO*INT_ (2, 85) = 3.18, *p* = 0.047).

**Figure 10.**
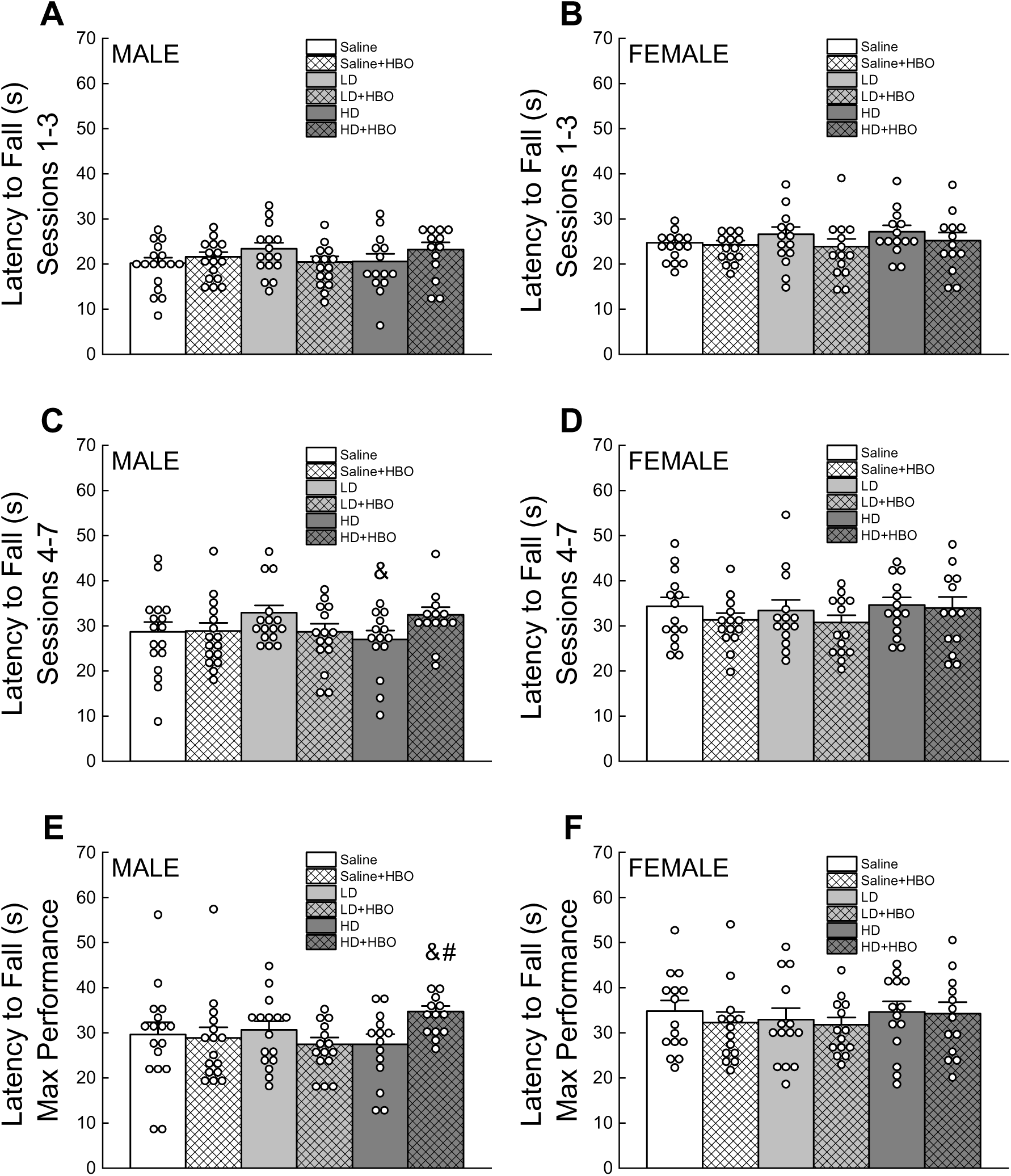
Effects of chemotherapy and HBO on coordinated running (rotorod). Bar graphs depict latency to fall during sessions 1-3 (**A**, **B**), sessions 4-7 (**C**, **D**), and last session (**E**, **F**) of male (left panels) and female (right panels) mice. Data are represented as mean ± S.E.M (Male: n=13-17; Female: n=13-15). & p<0.05 versus intervention-matched LD; # p<0.05 versus chemotherapy-matched non-HBO.

In females (Figs. 10A, 10C, 10E), there was no effect of chemotherapy or HBO on any of the measures of coordinated running (all Fs < 2.400, all *ps* > 0.130).

### 5.8 Anxiety-like behavior

In males (Figs. 11A, 11C), there was a trend of decreased distance for the LD and HD groups, however the main effect did not reach significance (F_CHEMO_ (2, 86) = 2.525, *p* = 0.086). There was no effect of chemotherapy or HBO on time spent in open arms (all Fs < 1.20, all *ps* > 0.30). In females (Figs. 11B, 11D), there was a trend of decreased distance for the HD groups, however the main effect did not reach significance (F_CHEMO_ (2, 74) = 2.892, *p* = 0.062). There was no effect of chemotherapy or HBO on time spent in open arms (all Fs < 1.72, all *ps* > 0.15).

**Figure 11.**
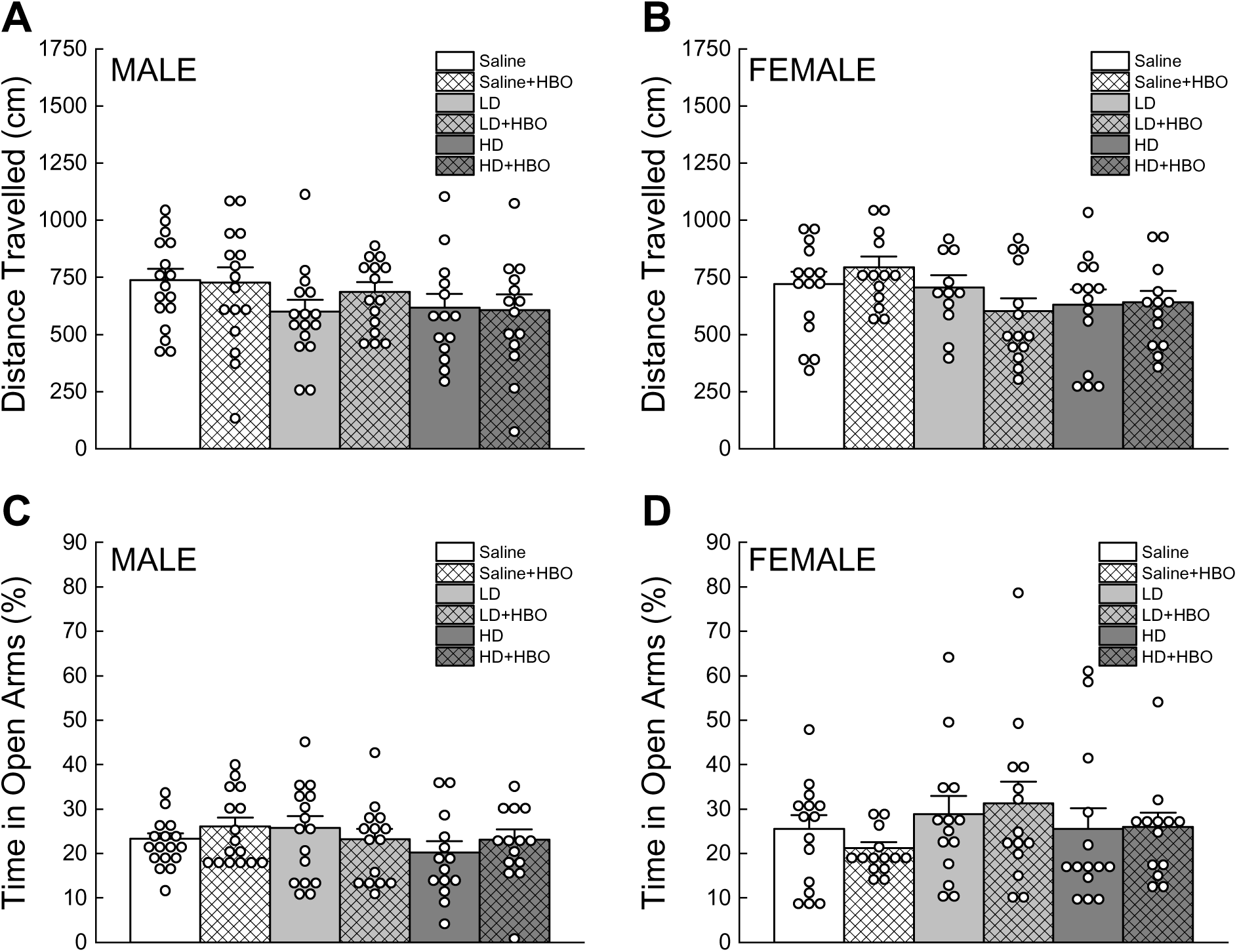
Effects of chemotherapy and HBO on anxiety-related behaviors. Bar graphs depict average distance travelled (**A**, **B**) and % time spent in the open arms (**C**, **D**) during a 5-minute test of male (left panels) and female (right panels) mice. Data are presented as mean ± S.E.M (Male: n=14-17; Female: n=11-15).

## 6. Discussion

This study examines the effects of HBO in a mouse model of CRCI, evaluating its impact on general health, cognitive, motor, and affective functions. Our findings suggest that HBO modulated chemotherapy-induced deficits in a dose- and task-dependent manner. (1) Chemotherapy, particularly at higher dose, impaired spatial memory, avoidance learning, contextual fear discrimination, spontaneous locomotion (exploration) and anxiety-like behavior in males, and (2) impaired spatial memory and avoidance learning in females. (3) HBO significantly improved avoidance learning in both sexes, however it also appeared to (4) worsen spatial memory deficits in males and anxiety in females. While chemotherapy did not affect motor function, (5) HBO seemed to improve maximal running performance of high-dose males.

Chemotherapy-induced toxicity is well-documented, leading to systemic effects such as weight loss, skeletal muscle wasting, fatigue, and hair loss [22, 73–77]. In line with those findings, low-dose (LD) temporarily induced hair loss and poor coat conditions in males, while high-dose (HD) temporarily decreased food intake in males and induced weight loss in both sexes, with recovery by the end of the study. Despite weight loss, no significant muscle weakness was observed, contrasting with a prior study [78]. The dose-dependent impact on hair and body weight could stem from metabolic differences and compensatory mechanisms, where LD disrupt the hair growth cycle without impacting body weight, whereas HD toxicity suppresses appetite, resulting in weight loss, but may trigger stronger compensatory pathways promoting faster hair regrowth [79]. However, all animals recovered after treatment cessation, suggesting transient effects of chemotherapy on these measures. Consistent with previous works showing repeated exposure to HBO increases muscle fiber growth and promotes muscle regeneration after contusion injury [80, 81], HBO mitigated weight loss and improved grip strength in females, but not males. This differential response may stem from females having higher muscle oxidative metabolism [82], which benefits more from the increased oxygen availability of HBO, thus improving muscle mass and weight recovery. Overall, our findings suggest that chemotherapy-induced toxicity is transient and that HBO dampened its impact and/or fastened recovery especially in females.

Even small doses of common chemotherapeutic agents can cause cell death and alter brain structures crucial for cognition [83]. Both MTX and 5-FU cross the blood-brain-barrier, potentially exerting direct neurotoxic effects [84, 85]. As expected, chemotherapy, especially HD, impaired spatial and associative avoidance learning and memory in both sexes, impaired spatial navigation and contextual discrimination in males. These deficits are consistent with previous CRCI models demonstrating impairment most pronounced in hippocampal and frontal lobe functions [30, 86–89]. Additionally, the hippocampal-amygdala circuit is involved in contextual and cued memory [71] with the amygdala associated with fear responses to both cue and context whereas hippocampus is associated with context [90]. We showed that males exposed to HD exhibited heightened freezing in the novel context but not in the presence of the conditioning stimulus, supporting the hypothesis that the hippocampus is more sensitive to chemotherapy compared to other brain regions [91]. Increased fear generalization may result from impaired safety signal learning and inability to suppress fear responses in safe environments [92, 93]. Males and females do respond differently to fear conditioning [94], and the lack of HD effect in females suggest a potential influence of sex hormones [94].

HBO exposure resulted in mixed outcomes on cognitive function. It improved discriminated avoidance learning in both sexes, consistent with prior studies in hypoxic ischemia and Alzheimer’s disease [95, 96]. However, HBO did not improve spatial learning and memory and may even impair function and delay use of spatial strategies in HD males, contradicting prior reports of HBO improving water maze performance in animal models of neurodegenerative diseases [97–100].

Cancer patients frequently experience affective disturbances, with males exhibiting greater levels of anxiety and females exhibiting higher levels of depression [101]. Reduced exploration and/or time spent in the open area could indicate increased anxiety [102, 103]. Chemotherapy increased anxiety-like behaviors in males, evidenced by reduced exploration and time spent in the center of open field consistent with previous findings [104], but did not affect performance in the elevated plus maze. This divergence between anxiety outcomes measured by these two different tests has been observed before [105]. HBO did not alleviate these symptoms, and, in females, appeared to increase anxiety when combined with chemotherapy. The findings align with prior research on the multidimensional structure of anxiety behaviors, which are sex- and task-dependent [106, 107]. The lack of therapeutic HBO effects on affective function is consistent with studies in AD model [96] but contrasts with its anxiolytic effects in spinal cord injury [108], COVID-19 [109], chronic stress [110], and chronic pain-related conditions [111, 112]. Chronic pain from central sensitization, where persistent pain leads to changes in the central nervous system amplifying pain perception, may result in emotional distress as secondary effects [113–116]. Thus, the anxiolytic effects of HBO in these conditions may be secondary as it provided significant chronic pain relief [117], however we did not measure pain indices in this study.

Chemotherapy had no observable effects on motor coordination of either sexes, consistent with prior findings [118]. Nonetheless, HBO enhanced maximum coordinated running capacity in HD-treated males, aligning with its established benefits in TBI, stroke, and neurodegenerative diseases [96, 119–122]. Previous studies suggested that HBO’s motor effects are task-dependent, selectively benefiting more challenging tasks. In males, pre-exposure to HBO improved performance on a larger diameter rotorod [123], while delayed exposure aided recovery in faster accelerating rotorod after ischemic stroke [124]. This suggests HBO is more effective under higher motor demands, potentially aiding recovery exclusively in more severely affected systems such as those treated with high-dose chemotherapy. Effects were also seemingly sex- and domain-specific as suggested previously [96], potentially due to sex differences in white matter neuroplasticity following motor/balance skill acquisition as males and females possibly rely on distinct brain regions to achieve motor outcomes [125].

While we aimed to assess effects of HBO on CRCI, the current chemotherapy regimen did not consistently produce cognitive impairments across functional domains. This inconsistency contrasts with prior studies using comparable LD chemotherapy models [30, 86, 87], and could be due to strain differences. A lack of robust effects on some measurements may also be due to the fact that CRCI are not persistent in mouse models (some of the behavioral tests were done 2- 3 weeks after the last chemo injections). Use of immunocompromised mouse model or a model with induced cancer or more prolonged exposure to chemotherapy may have led to more significant impairments of chemotherapy. A more robust CRCI model may be necessary to further evaluate HBO as a viable intervention. While it seems that HBO was able to improve some cognitive domains significantly, others had small effects. HBO’s regimen was chosen based on previous work in our lab in an AD mouse model [96], but frequency and duration of HBO exposure might need to be optimized for different conditions.

## 7. Conclusions

Our findings add to the literature supporting HBO as a therapy for conditions affecting brain function including CRCI. The complexity of CRCI and underlying mechanisms may explain the nuance and mixed outcomes of HBO on different domains. Our study also highlights the importance of the consideration of sex in CRCI and intervention research, as males and females responded differently to the chemotherapeutic treatment as well as to HBO exposure. Future work is necessary to further study the underlying mechanisms of the effects of HBO in both sexes, as well as refining HBO treatment specifics to maximize its benefits, including its long-term effects and duration of beneficial effects.

## 8. Credit authorship contributions

**Oanh T.P. Trinh**: Conceptualization, Methodology, Investigation, Data curation, Formal analysis, Visualization, Writing – original draft, Writing – review & editing, Funding acquisition. **Andrew Ferns**: Software (for MWM search strategies). **Bethany Zachariah**: Data curation. **Nathalie Sumien**: Conceptualization, Methodology, Data curation, Formal analysis, Writing – review & editing, Resources, Supervision, Project administration, Funding acquisition.

## Acknowledgements

This study was sponsored by The Feddersen Trust Foundation and the Lowdon Family Foundation. Author Oanh T.P. Trinh was partially supported by a grant from the Cancer Prevention and Research Institute of Texas (Award#: RP210046) to Dr. Jamboor Vishwanatha.

---

**Supplementary Figure 1.**
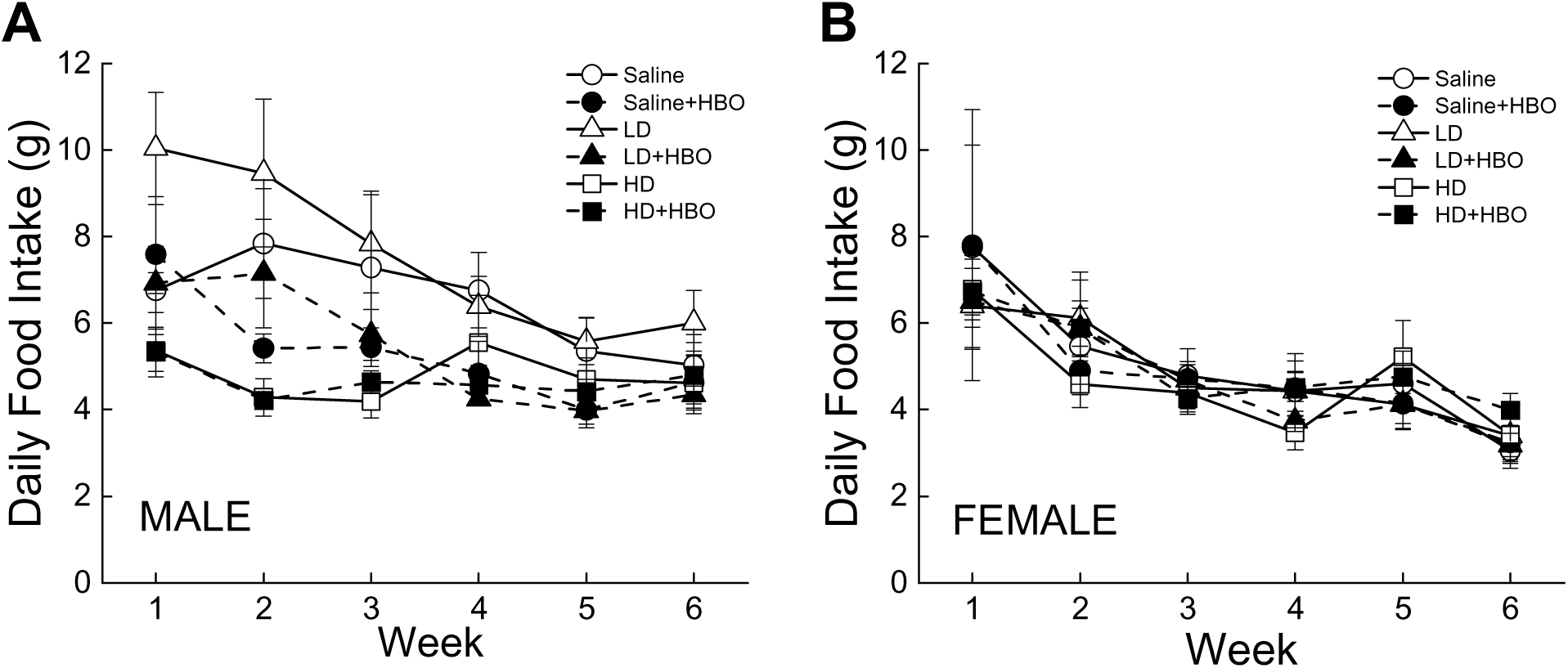
Effects of chemotherapy and HBO on daily food intake. Line graphs depict the daily average food consumption of male (**A**) and female (**B**) mice. Data are presented mean ± S.E.M (Number of cages: Male: n=3-5; Female: n=3-5).

**Supplementary Figure 2.**
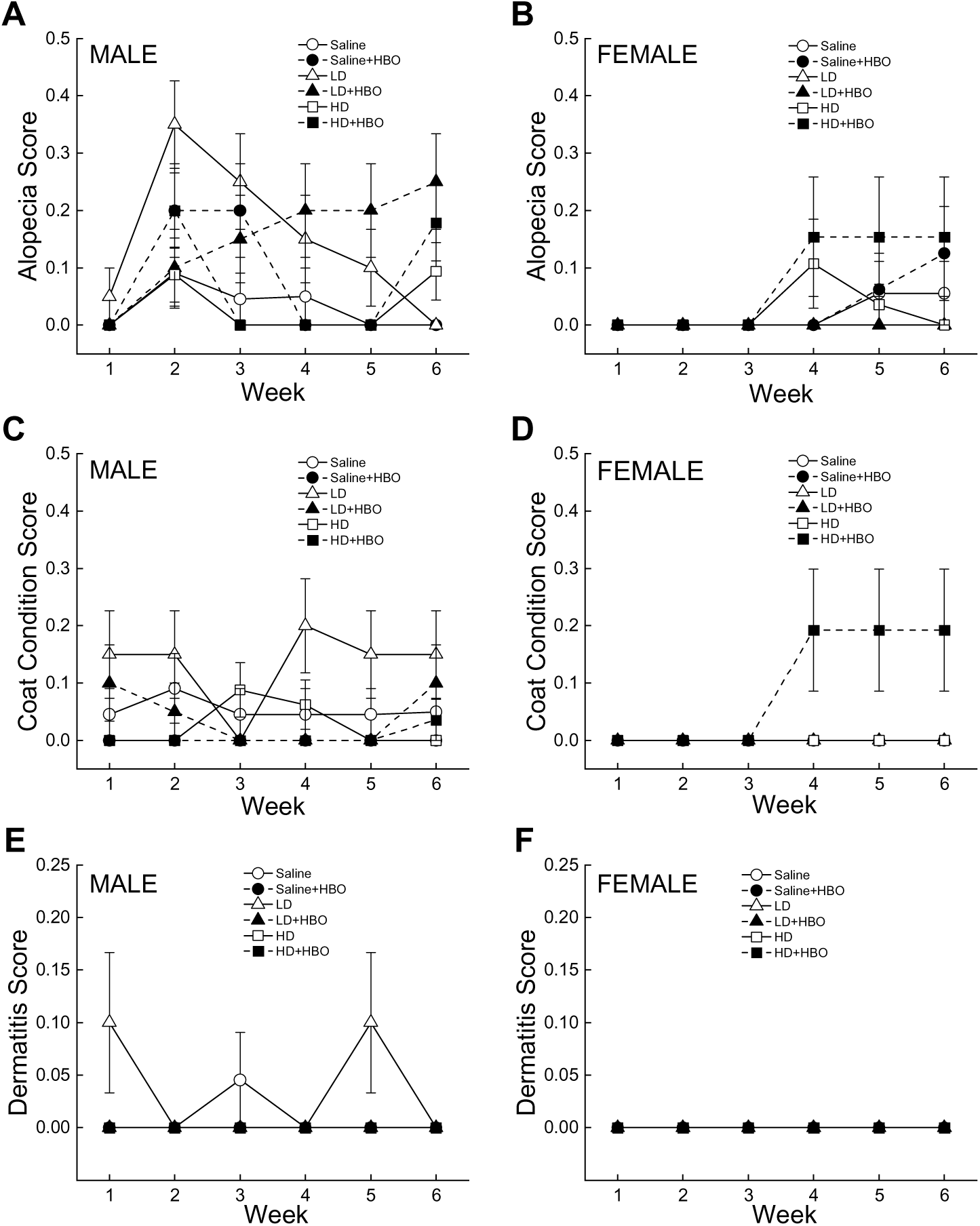
Effects of chemotherapy and HBO on clinical observations. Line graphs depict the weekly average scores on alopecia (**A**, **B**), coat condition (**C**, **D**), and dermatitis/skin condition (**E**, **F**) of male (left panels) and female (right panels) mice. Clinical conditions were evaluated according to the following scale: 0 = absent, 0.5 = mild, 1 = severe. Data are presented mean ± S.E.M (Male: n=3-5; Female: n=3-5).

**Supplementary Table 1.** Percentage of search strategies during Morris water maze.

| STRATEGY | MALE |  |  |  |  |  |  |  | FEMALE |  |  |  |  |  |  |  |
| --- | --- | --- | --- | --- | --- | --- | --- | --- | --- | --- | --- | --- | --- | --- | --- | --- |
|  | Session 1 |  | Session 2 |  | Session 3 |  | Session 4 |  | Session 1 |  | Session 2 |  | Session 3 |  | Session 4 |  |
| Saline | — | +HBO | — | +HBO | — | +HBO | — | +HBO | — | +HBO | — | +HBO | — | +HBO | — | +HBO |
| Direct Path | 0.0 | 1.3 | 5.9 | 4.7 | 10.6 | 7.5 | 5.9 | 6.3 | 0.0 | 2.9 | 5.0 | 8.9 | 5.3 | 14.3 | 11.7 | 7.1 |
| Directed Search | 0.0 | 0.0 | 10.3 | 4.7 | 5.9 | 15.0 | 14.7 | 14.1 | 8.0 | 0.0 | 10.0 | 1.8 | 17.3 | 14.3 | 13.3 | 3.6 |
| Focal Search | 2.4 | 0.0 | 2.9 | 0.0 | 4.7 | 6.3 | 5.9 | 4.7 | 0.0 | 0.0 | 5.0 | 3.6 | 8.0 | 2.9 | 8.3 | 10.7 |
| Indirect Search | 7.1 | 10.0 | 23.5 | 34.4 | 37.6 | 27.5 | 25.0 | 32.8 | 17.3 | 17.1 | 33.3 | 35.7 | 25.3 | 25.7 | 21.7 | 35.7 |
| Chaining | 0.0 | 0.0 | 0.0 | 0.0 | 0.0 | 0.0 | 0.0 | 0.0 | 0.0 | 0.0 | 0.0 | 0.0 | 0.0 | 0.0 | 0.0 | 0.0 |
| Scanning | 11.9 | 12.5 | 14.7 | 10.9 | 20.0 | 8.8 | 17.6 | 10.9 | 9.3 | 15.7 | 15.0 | 7.1 | 16.0 | 8.6 | 15.0 | 14.3 |
| Random Search | 78.7 | 76.3 | 42.6 | 45.3 | 21.2 | 35.0 | 30.9 | 31.3 | 65.3 | 64.3 | 31.7 | 42.9 | 28.0 | 34.3 | 30.0 | 28.6 |
| Thigmotaxis | 0.0 | 0.0 | 0.0 | 0.0 | 0.0 | 0.0 | 0.0 | 0.0 | 0.0 | 0.0 | 0.0 | 0.0 | 0.0 | 0.0 | 0.0 | 0.0 |
|  | Session 1 |  | Session 2 |  | Session 3 |  | Session 4 |  | Session 1 |  | Session 2 |  | Session 3 |  | Session 4 |  |
| Low-dose | — | +HBO | — | +HBO | — | +HBO | — | +HBO | — | +HBO | — | +HBO | — | +HBO | — | +HBO |
| Direct Path | 1.3 | 0.0 | 4.7 | 5.0 | 6.3 | 6.7 | 14.1 | 21.7 | 1.4 | 1.4 | 7.1 | 5.4 | 7.1 | 11.8 | 14.3 | 8.9 |
| Directed Search | 2.5 | 5.3 | 7.8 | 6.7 | 12.5 | 12.0 | 12.5 | 16.7 | 4.3 | 5.7 | 3.6 | 3.6 | 12.9 | 8.7 | 14.3 | 16.1 |
| Focal Search | 0.0 | 0.0 | 1.6 | 1.7 | 6.3 | 5.3 | 10.9 | 8.3 | 0.0 | 1.4 | 3.6 | 3.6 | 10.0 | 0.0 | 14.3 | 5.4 |
| Indirect Search | 12.5 | 13.3 | 25.0 | 26.7 | 30.0 | 29.3 | 25.0 | 23.3 | 8.6 | 15.7 | 44.6 | 32.1 | 34.3 | 18.8 | 21.4 | 17.9 |
| Chaining | 0.0 | 0.0 | 0.0 | 0.0 | 0.0 | 0.0 | 0.0 | 0.0 | 0.0 | 0.0 | 0.0 | 0.0 | 0.0 | 0.0 | 0.0 | 0.0 |
| Scanning | 11.3 | 8.0 | 12.5 | 21.7 | 10.0 | 20.0 | 9.4 | 11.7 | 12.9 | 14.3 | 7.1 | 14.3 | 8.6 | 10.3 | 8.9 | 5.4 |
| Random Search | 72.5 | 73.3 | 48.4 | 38.3 | 35.0 | 26.7 | 28.1 | 18.3 | 72.9 | 61.4 | 33.9 | 41.1 | 27.1 | 50.4 | 26.8 | 46.4 |
| Thigmotaxis | 0.0 | 0.0 | 0.0 | 0.0 | 0.0 | 0.0 | 0.0 | 0.0 | 0.0 | 0.0 | 0.0 | 0.0 | 0.0 | 0.0 | 0.0 | 0.0 |
|  | Session 1 |  | Session 2 |  | Session 3 |  | Session 4 |  | Session 1 |  | Session 2 |  | Session 3 |  | Session 4 |  |
| High-dose | — | +HBO | — | +HBO | — | +HBO | — | +HBO | — | +HBO | — | +HBO | — | +HBO | — | +HBO |
| Direct Path | 1.4 | 0.0 | 1.8 | 1.9 | 0.0 | 7.7 | 7.1 | 5.8 | 0.0 | 0.0 | 5.0 | 7.7 | 5.8 | 9.2 | 7.1 | 7.7 |
| Directed Search | 2.9 | 0.0 | 1.8 | 5.8 | 2.9 | 4.6 | 10.7 | 9.6 | 0.0 | 6.2 | 5.0 | 5.8 | 13.0 | 7.7 | 10.7 | 17.3 |
| Focal Search | 0.0 | 0.0 | 0.0 | 0.0 | 1.4 | 3.1 | 1.8 | 7.7 | 0.0 | 0.0 | 1.7 | 3.8 | 3.0 | 6.2 | 1.8 | 5.8 |
| Indirect Search | 11.4 | 7.7 | 16.1 | 13.5 | 22.9 | 26.2 | 23.2 | 28.8 | 14.7 | 13.8 | 25.7 | 13.5 | 23.1 | 16.9 | 28.6 | 19.2 |
| Chaining | 0.0 | 0.0 | 0.0 | 0.0 | 0.0 | 0.0 | 0.0 | 0.0 | 0.0 | 0.0 | 0.0 | 0.0 | 0.0 | 0.0 | 0.0 | 0.0 |
| Scanning | 14.3 | 1.5 | 28.6 | 5.8 | 30.0 | 15.4 | 26.8 | 9.6 | 9.3 | 12.3 | 10.2 | 25.0 | 19.0 | 20.0 | 10.7 | 19.2 |
| Random Search | 70.0 | 90.8 | 51.8 | 73.1 | 42.9 | 43.1 | 30.4 | 38.5 | 76.0 | 67.7 | 52.4 | 44.2 | 36.2 | 40.0 | 41.1 | 30.8 |
| Thigmotaxis | 0.0 | 0.0 | 0.0 | 0.0 | 0.0 | 0.0 | 0.0 | 0.0 | 0.0 | 0.0 | 0.0 | 0.0 | 0.0 | 0.0 | 0.0 | 0.0 |

